# Akaike information criteria and predictive geographical accuracy are not related in ecological niche modeling

**DOI:** 10.1101/315101

**Authors:** Julián A. Velasco, Constantino Gonzales-Salazar

**Affiliations:** Museo ‘Alfonso L. Herrera’, Facultad de Ciencias, Universidad Nacional Autónoma de México, Mexico city, Mexico.; Centro de Ciencias de la Complejidad –C3- Universidad Nacional Autónoma de México, Mexico city, Mexico.

**Keywords:** Distribution, Ecological niche, Geography, Maxent, Validation, Virtual species

## Abstract

**Aim:** Ecological niche modeling (ENM) is an approach used to estimate species‘ presence given its environmental preferences. Model complexity in ENMs has increasingly gained relevance in the last years. In particular, in Maxent algorithm is captured using the Akaike Information Criteria (AIC) based on the number of parameters and likelihoods of continuous raw outputs. However, it is not clear whether best-selected models using AIC are the models with the highest classification rate of correct presences and absences. Here, we test for a link between model complexity and accuracy of geographical predictions of Maxent models.

**Innovation:** We created a set of virtual species and generate true geographical predictions for each one. We build a set of Maxent models using presence data from each virtual species with different regularization and features schemes. We compared AICc values for each model with the scores of standard validation metrics (e.g., Kappa, TSS) and with the number of pixels correctly predicted as presences, absences or both.

**Main Conclusions:** We found that binary predictions (i.e., presence-absence maps) selected as best models for AIC tend to predict incorrectly sites as presences and absences using independent datasets. We suggest that information criteria as AIC should be avoided when users are interested in binary predictions. Future applications that capture model complexity in ENM applications should be evaluated using standard validation metrics.

## Introduction

Ecological niche modeling (henceforth ENM) is a conceptual approach that allows estimating where a species occurs in a region based on a set of coarse-grain environmental variables (Peterson et al., 2011, Franklin, 2010). This approach is grounded firmly in ecological niche theory and to estimate, at least theoretically, where species can have positive intrinsic growth rates (i.e., source populations; Maguire, 1973; Pulliam, 2000; Holt, 2009; Soberón, 2010; Peterson et al., 2011). ENM approaches are used for multiple purposes including, but not limited, estimates of potential and realized geographical distributions, range shifts due to environmental change, or predictions of sites with the high potential risk of invasive species or disease outbreaks (Peterson et al., 2011; Araújo & Peterson, 2012).

ENM approaches use a mathematical algorithm (e.g., machine learning, spline regressions) to establish a link between species’ presence data and environmental layers across a geographic region (Merow et al., 2013). The environmental data for species occurrences are extracted from a geographical space and a model is fitted in an *n*-dimensional space (i.e., the ecological space). This fitted model is projected to a geographical region to predict where a species can have a high probability of occurrence given its environmental preferences (Peterson et al., 2011; Peterson & Soberón, 2012). Several types of mathematical algorithms are used for ENMs depending on the availability of species data. Some techniques only use presence data (e.g., Bioclim), whereas others use presence and absence data (e.g., GLM, GAM). However, as absence data are very hard to collect (Peterson et al., 2011, some techniques to generate a random sample of points across the study region to be used as pseudo-absence data (e.g., GARP; Stockwell, 1999) or background data (e.g., Maxent; Phillips & Dudík, 2008).

The model outputs are taken as geographical predictions that can be considered, given a specific set of assumptions, as the probability of presence (Phillips & Dudik, 2008; but see Merow et al., 2013) or suitability values (Merow et al., 2013). Usually, a thresholding is implemented to convert continuous outputs in binary predictions (i.e., presence-absence data) using a particular criterion (e.g., minimum training presence; Liu et al., 2005). Therefore, binary model outputs are simply geographical predictions of where species can be present or absent given its ecological niche preferences. Thresholding usually is employed by users interested in prediction-oriented models (Peterson et al., 2011). These binary predictions are validated with an independent set of presences and absences (or pseudo-absences) data (Fielding & Bell, 1997). Model validation is conducted in an explicit geographical context to establish whether presence and absence sites are predicted correctly using different types of metrics (Fielding & Bell, 1997; Allouche et al., 2006; Peterson et al., 2011).

Recent progress in ENMs has involved the notion of model complexity and information contended in models (Segurado & Araújo, 2004; Elith et al., 2006; Warren & Seinfert, 2011; Warren et al., 2014; García-Callejas & Araújo, 2015). Warren & Seinfert (2011) implemented the Akaike Information Criteria (AIC; Anderson & Burham, 2002) to choose the best models based on the number of parameters and model likelihoods for the Maxent algorithm. With this approach, it is possible to select the best model between different settings of model parameters changing features and regularization values. However, still is uncertain whether best models selected with AIC tend to minimize the error in predicted presences and absences in binary outputs. As binary outputs are validated in an explicit geographical context, it is necessary to establish whether best models selected with AIC are those with the highest predictive power in the geographical context.

In this paper, we ask whether optimal model complexity, captured through AIC (Warren & Seinfert, 2011; Muscarella et al., 2014), is linked with the ability of models to discriminate well between presences and absences in an explicit geographical context. We expect that models with lower AICc values (i.e., the best models) tend to predict correctly an independent dataset of presence and absences. By contrast, models with higher AICc values (i.e., the worst models) are expected to be unable to discriminate correctly between presence and absence data. However, our results suggest that Akaike Information Criteria is not correlated with several validation metrics and therefore should not be used when researchers are interested in a prediction-oriented model, particularly a presence-absence prediction. We recommend that new applications oriented to measure model complexity also should be tested in a geographical context using standard validation metrics (e.g., Kappa, TSS).

## Methods

### Generation of virtual species and true maps of presence and absence

We generated three virtual species based on the distribution of three common mammals distributed in North America (Table 1). Virtual species allow us to establish correctly where a species can be found (i.e., true presences) and where is not (i.e., true absences) (Miller, 2014). The ecological niche of each species was defined as ellipsoid from three climatic variables: annual mean temperature (bio1), annual precipitation (bio12) and precipitation seasonality (bio15) from the WorldClim database (Hijmans et al., 2005). We assumed that fundamental niches exhibit an ellipsoid shape in the multivariate ecological space (Soberón & Nakamura, 2009; Soberón, J. pers. comm.). We used a minimum volume ellipsoid model to estimate the fundamental niche of each virtual species. This kind of model uses a rescaled multinormal probability density function (Osorio-Olvera et al., 2016a) to generate a suitability index which is reflected in the geographical space (Figure 1). We used the NicheToolBox package (Osorio-Olvera et al., 2016b) available in the GitHub repository (https://github.com/luismurao/nichetoolbox) to generate suitability raster maps for each species. Each suitability map was reclassified to generate a binary prediction of true presence and absence of each virtual species in the geography. All suitability values equal to zero were considered as absences and all suitability values greater than zero were considered as presences. From each true presence-absence map, we extracted a random sample of pixels predicted as presences and absences for each virtual species (Table 1). The presences were randomly split to generate training and testing datasets which were used to calibrate and validate niche models. The number of absences for validating each model was calculated based on the prevalence of each virtual species (Table 1).

**Table 1.** Virtual species used in this study based on the distribution of three North American mammals. Threshold value refers to the value used to generate a binary (presence-absence) geographical predictions from continuous outputs. The number of presence and absence records used to calibrate and validate models, respectively.

| Virtual species | Threshold value | Training presence records | Testing absences records | Known mammal species |
| --- | --- | --- | --- | --- |
| Species 1 | 0.010 | 203 | 1822 | *Aplodontia rufa* |
| Species 2 | 0.010 | 219 | 1809 | *Blarina carolinensis* |
| Species 3 | 0.010 | 133 | 1883 | *Blarina hylophaga* |

**Fig. 1.**
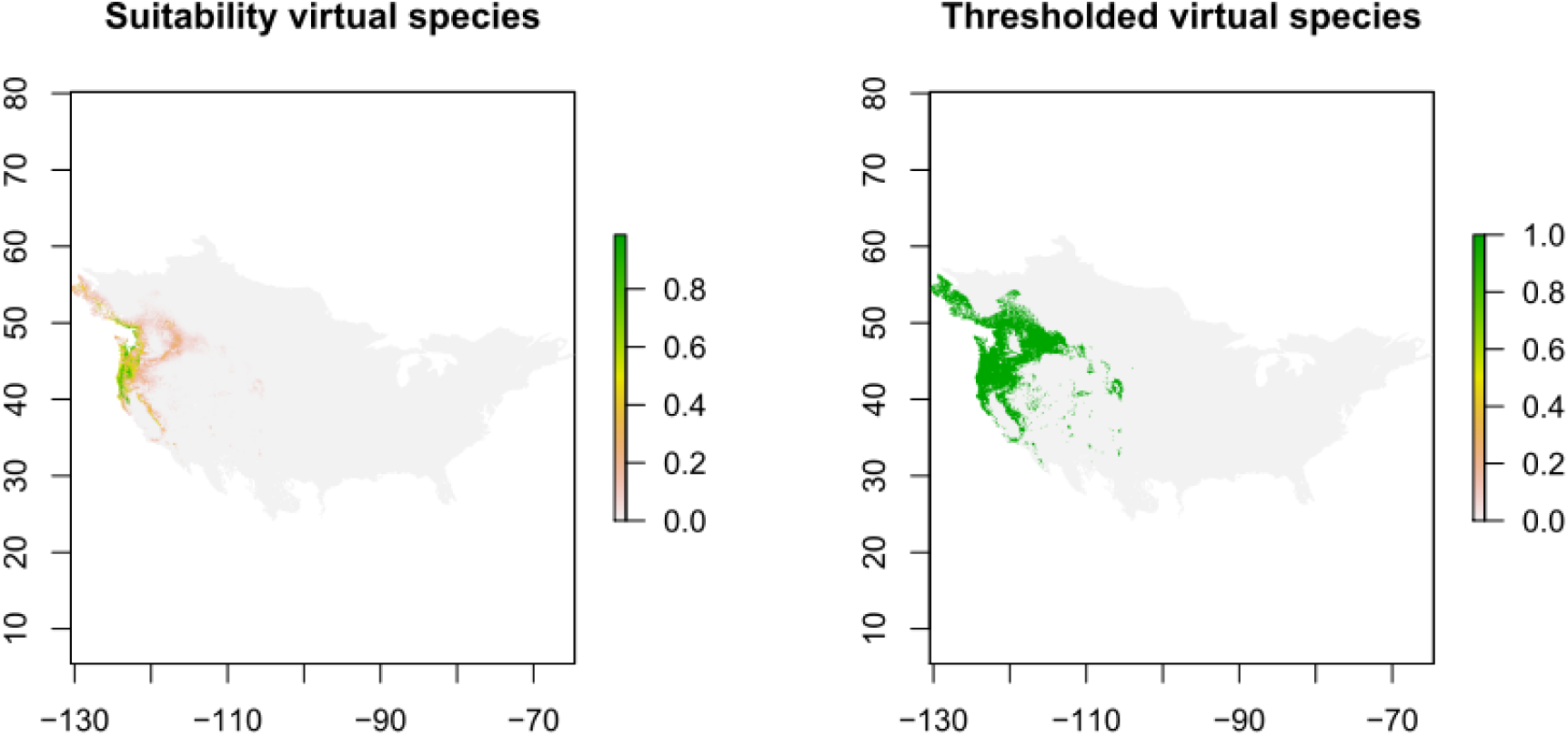
Visualization of the ecological niche of one of the virtual species used in this study in both ecological space (E-space) and geographical space (G-space) in the study area.

### Ecological niche modeling, model parameterization

Using a presence training dataset for each virtual species we generated Maxent models with different regularization values and feature classes using the presences training dataset for each species (Table 1). The regularization is a function implemented to avoid overfitting in models (Phillips et al., 2006) and as this value increases the model predictions tend to sparse out in the geography (Phillips et al., 2006; Radosavljevic & Anderson, 2014). The feature classes are functions that link variables with species’ presences, some are very simple (e.g., linear) whereas others are very complex (e.g., hinge or threshold; Phillips & Dudik, 2008). We used the ENMevaluate function from the ENMeval R package (Muscarella et al., 2014) to generate a suite of models with different combinations of regularization values and features classes (H: hinge, L: linear, Q: quadratic, P: product and T: threshold). The regularization values ranged from 0.5 to 4 with increments of 0.5 and combinations of feature classes were as follows: L, LQ, H, LQH, LQHP, and LQHPT. For each species, we built 48 models and we calculated the AICc, delta AICc and weights ACIc based on the number of parameters used and the likelihood of each model. Models were built using a splitting-method based on jackknifing with 10 replicates. R scripts for generation of virtual species and ecological niche modeling are available in Appendix 1.

The AIC is an information metric that reflects both model goodness-of-fit and complexity (Burnham & Anderson, 2002; Warren & Seifert, 2011; Muscarella et al., 2014; Warren et al., 2014). The number of parameters from each model was counted as all parameters with a nonzero weight in lambdas (Warren & Seinfert, 2011). The log-likelihood is calculated as the product of the raw output scores for all pixels with a presence training record (Warren & Seinfert, 2011; see also Muscarella et al., 2014). The model with the lowest AICc is considered as the best model between all models. A thumb rule is usually used where models with delta AICc lower than 2 are considered as informative models (Burnham & Anderson, 2002).

### Model accuracy in binary predictions

To evaluate the effect of threshold selection in binary predictions, and therefore in the accuracy of prediction-oriented models, we reclassified models using several criteria for each virtual species (Liu et al., 2005; Peterson et al., 2011). We used the following criteria: minimum training presence (MTP), maximizing kappa (max.Kappa), maximizing sensitivity plus specificity (max.Sens.Spec), equal specificity and sensitivity (eq.Spec.Sens) and the same used in the virtual species (Same.Virtual; Table 1).

Each binary map was validated using an independent dataset of presences and absences (Table 1). The number of absences used to validate models was calculated based on the prevalence of the virtual species and the number of presences left to validate the model (Table 1). We calculated Kappa, True Skill Statistics (TSS), correct classification rate (CCR), incorrect classification rate (ICR), negative prediction power (NPP), positive prediction power (PPP) for each estimated binary map. We used this threshold-dependent validation metrics because our aim is to estimate the accuracy of prediction-oriented models instead of using threshold-independent metrics as AUC (Lobo et al., 2008). These validation metrics are calculated from the confusion matrix based on the presences and absences not used for calibration model (Fielding & Bell, 1997; Allouche et al., 2006; Peterson et al., 2011).

Kappa measures the accuracy of the binary model taking into account the ratio of correct predictions of presences and absences, although this metric is sensitive to the commission error (i.e., absences predicted incorrectly as presences). ENMs with Kappa values higher than 0.75 are considered as excellent models. True Skill Statistic (TSS) is the difference between the omission and commission errors. TSS ranges from -1 to 1, where zero indicates that the model is unable to differentiate between omission (presences predicted as absences) and commission (absences predicted as presences) errors, values close to 1 indicates that the model has a high predictive accuracy, and values close to -1 indicates a low predictive accuracy. Correct classification rate (CCR) measures the ratio of presences and absences points predicted correctly. Incorrect classification rate (ICR) measures the ratio of presences and absences points predicted incorrectly. Negative prediction power (NPP) evaluates the probability that true absence is correctly predicted as an absence. Positive prediction power (PPP) evaluates the probability that a true presence is correctly well predicted as presence (Fielding & Bell, 1997; Allouche et al., 2006; Peterson et al., 2011). Furthermore, we calculated the percentage of true presences (true positives), true absences (true negatives) and global (true positives and negatives) predictions comparing each estimated binary model with the true binary model (presence-absence of the virtual species). This calculation was performed overlapping the true and estimated binary rasters and count the number of pixels labeled as presences, absences, or global that were correctly predicted. Finally, we also evaluated how different sizes of training dataset (100, 75, 50, 25, 15, 10 and 5 training occurrence records) can affect the AIC scores and validation metrics. We performed this analysis using presence data for a single virtual species.

We evaluate how AIC values (AICc, delta AICc and Akaike weights) are related to the validation metrics capturing geographical prediction accuracy: Basically, we plotted the AICc values against each one of the validation metrics (i.e, AUC, TSS, Kappa, % true absences, % true presences, % true absences plus true presences). We expect to find a positive correlation between AIC scores and validation scores if both metrics are related. In other words, we expect to find that the highest scores of validation metrics are associated with the best-selected models using AIC.

## Results

Our results show that models with a poor performance based on AIC tend to exhibit a highest predictive accuracy using any standard metric of validation (Figure 2 and 3). We did not find a correlation between delta AICc values and different validation metrics of predictive accuracy model (Figure 2). ENMs obtain higher scores with multiple metrics independently of the AICc value (Figure 2). These results were similar independently of the threshold criteria used to generate binary map predictions (Figures S1-S6). We obtain similar results when we estimated the proportion of true presences and absences predicted correctly from “true” and estimated models (Figure 3; S7-S9). Although many models exhibited a high sensitivity (i.e., a higher number of presences predicted correctly) independently from delta AICc value (Figure 2, middle), by contrast, the specificity (i.e., the number of absences predicted correctly) was very variable across delta AICc values (Figure 2, top). These results were consistent across several threshold criteria used, but with higher variation for two criteria as minimum training presence and the same threshold value (Figure S7-S9). It seems that many models independently from its complexity (i.e., delta AICc value) predicted with high accuracy the sites where the species was present but failed in predict the sites where the species was absent. Some optimal models selected performed very poorly with different validations metrics (Figure 2 and 3). Even, we found that scores of validation metrics varied relatively across different regularization values and features schemes (Figure S10-S11). Finally, we showed that delta AIC values seem not be affected by sample size (Figure 4), but validation metrics (e.g., Kappa) were affected as the sample size decreased (Figure 5).

**Fig. 2.**
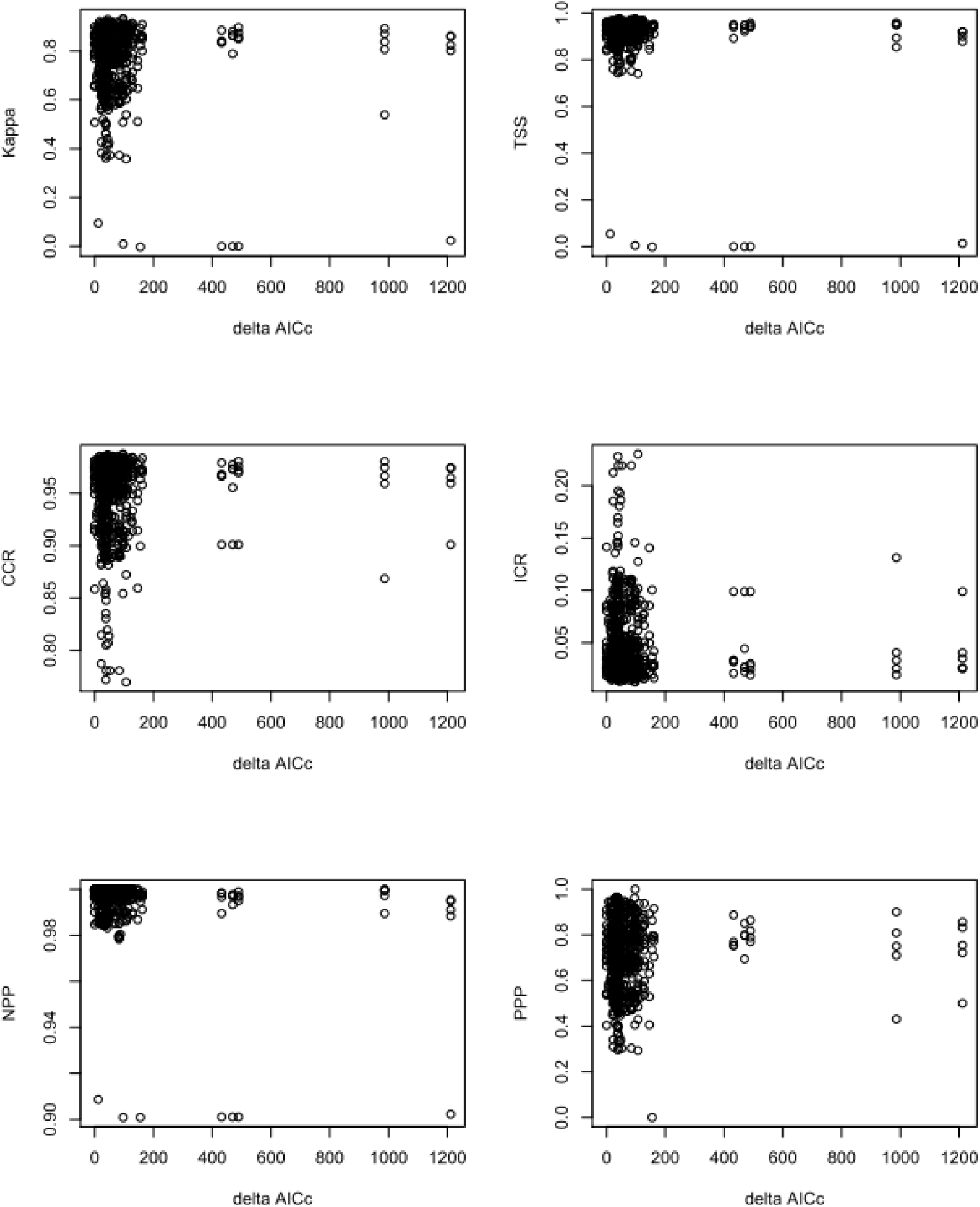
Plots between standard validation metrics of ecological niche models (ENMs) and delta AICc values. Delta AICc values represent the difference between a particular model with respect to the optimal (“best”) model. Accordingly, models with zero delta values and less than four represent the “most” optimal models based on its complexity (Burnham & Anderson 2002). TSS: true statistic skill; CCR: correct classification rate; ICR: incorrect classification; NPP: negative predictive power; PPP: positive predictive power (see main text for details of each metric).

**Fig. 3.**
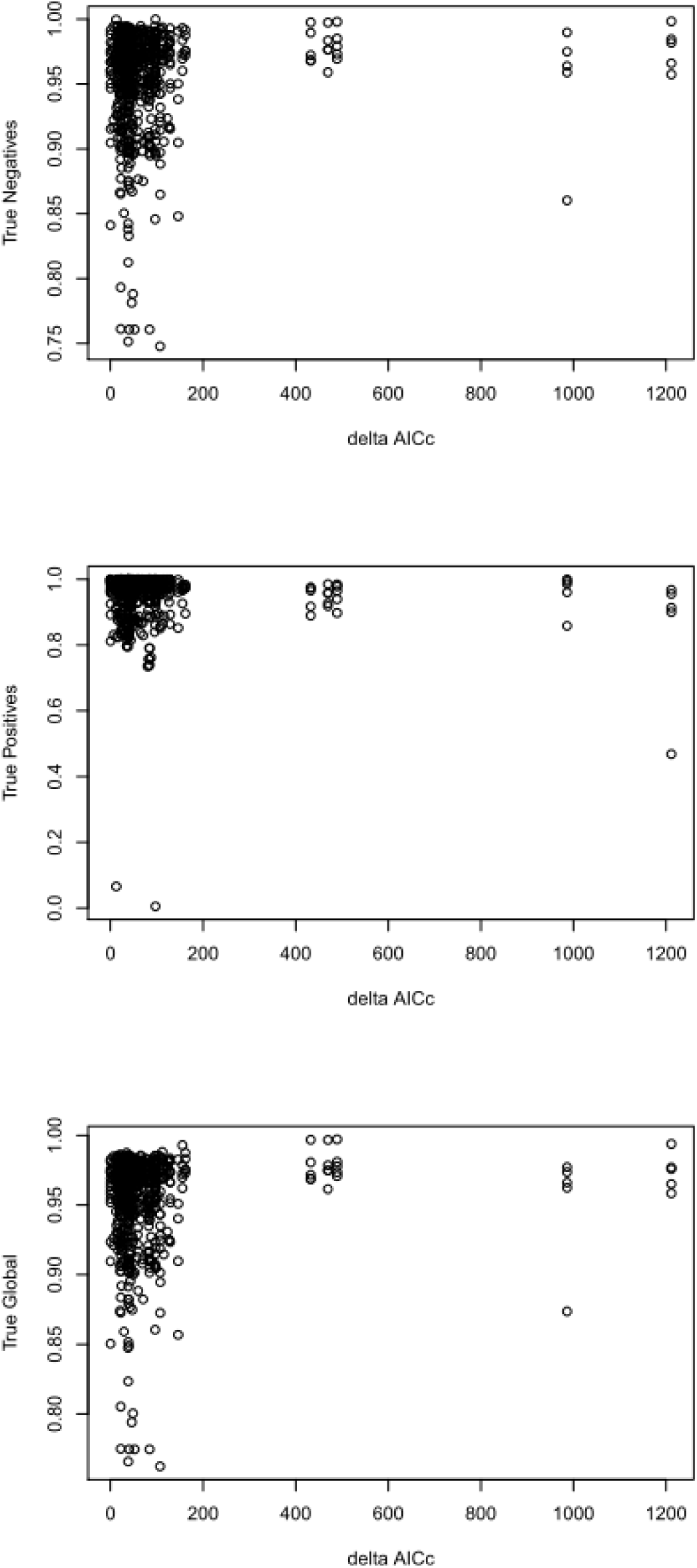
Plots of the percentage of true pixels correctly predicted from ecological niche models (ENMs) against delta AICc values. True negatives refer to the percentage of pixels labeled as absences in the true model and that was correctly predicted as absences in the estimated models. True positives refer to the percentage of pixels labeled as presences in the true model and that was correctly predicted as presences in the estimated models. True global refers to the percentage of true presences and absences combined in the true model and that was correctly predicted in the estimated models.

**Fig. 4.**
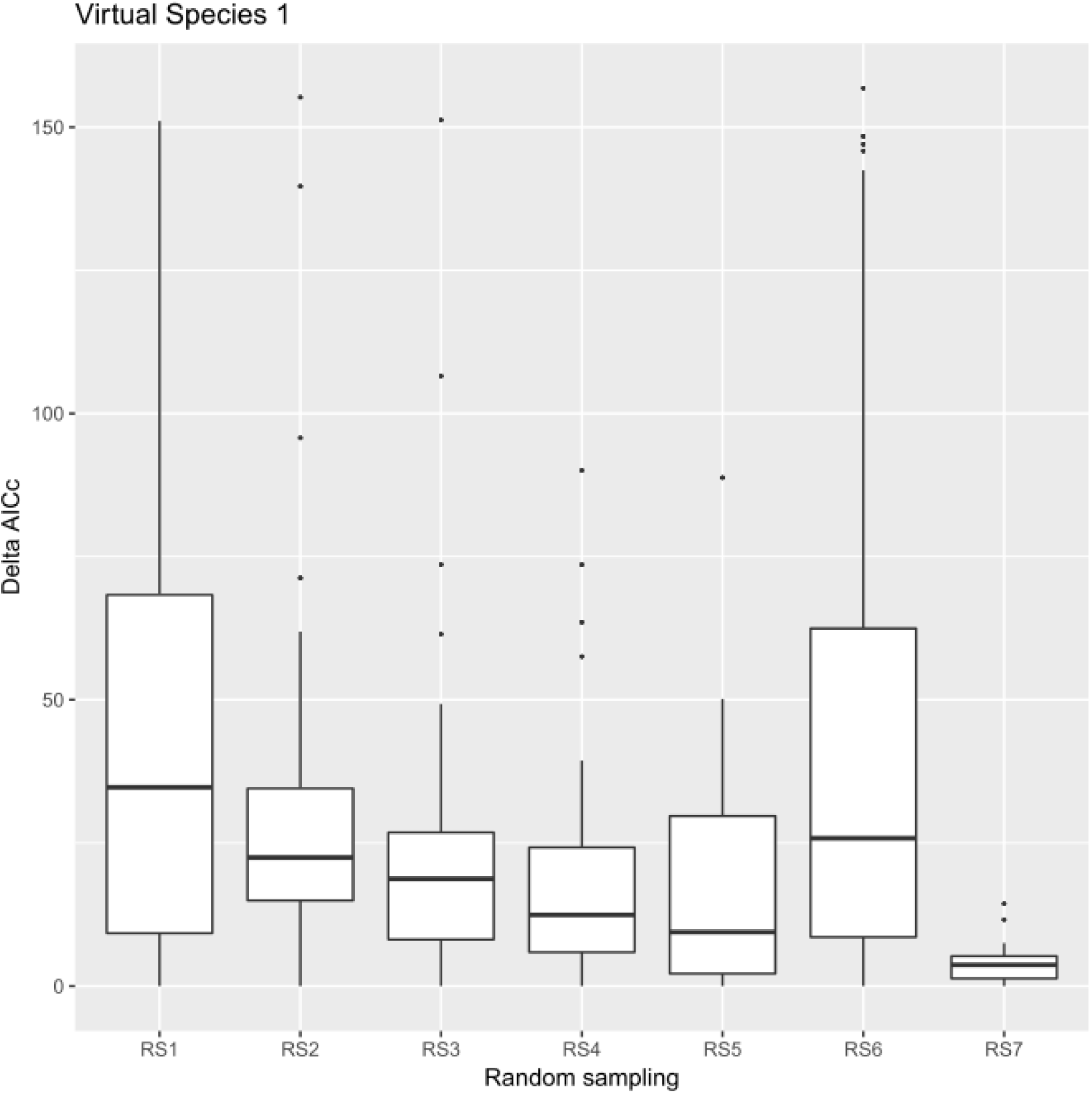
Boxplot of delta AICc values across models for different sample sizes in a training dataset (RS1 = 100 points RS2 = 75; RS3 = 50; RS4 = 25; RS5 = 15; RS6 = 10 and RS7 = 5).

**Fig. 5.**
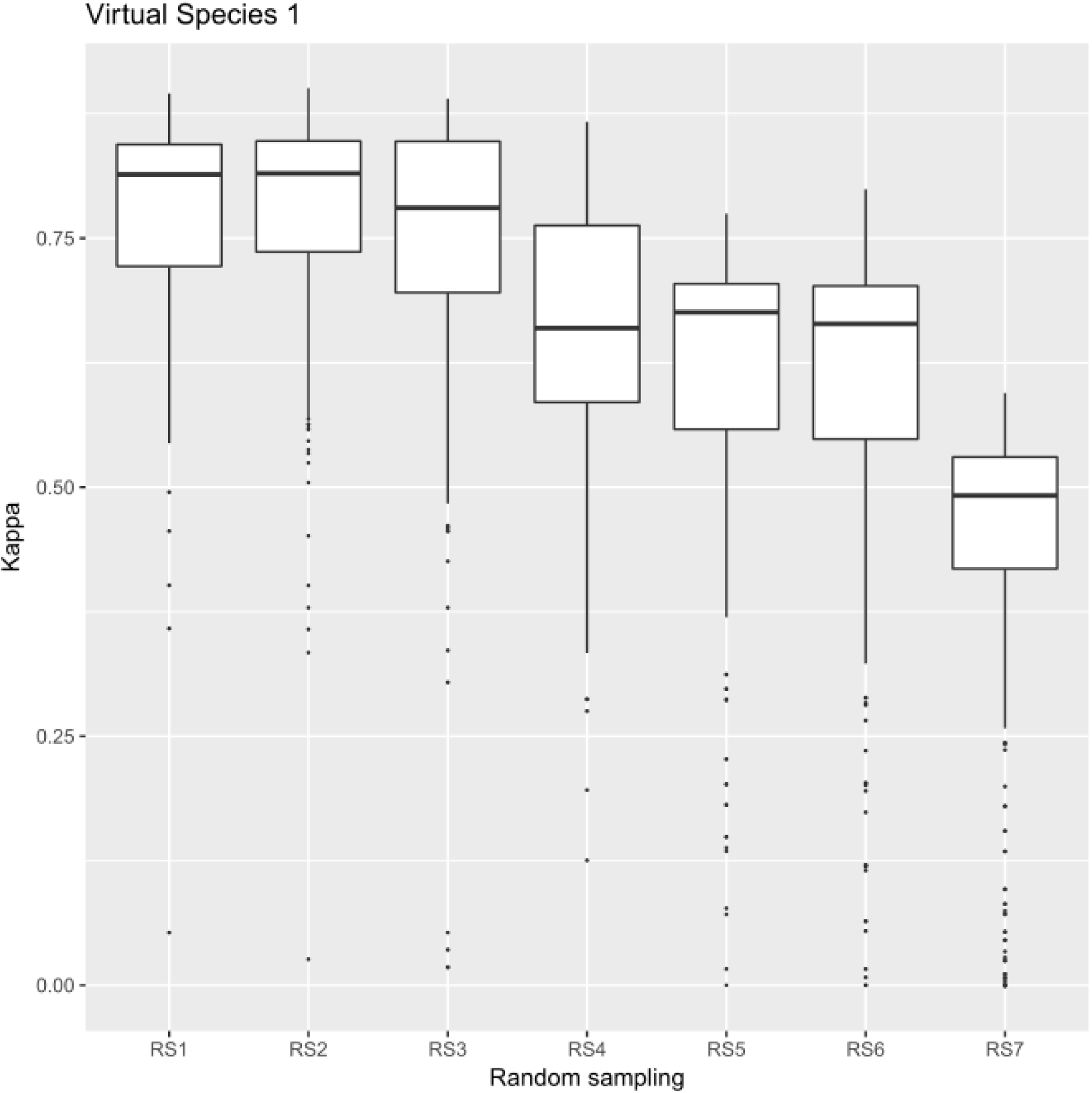
Boxplot of Kappa values across models for different sample sizes in a training dataset (RS1 = 100 points RS2 = 75; RS3 = 50; RS4 = 25; RS5 = 15; RS6 = 10 and RS7 = 5).

## Discussion

Previous studies have provided a link between ecological model complexity and predictive accuracy of ecological niche models (Warren & Seinfert, 2011; Warren et al., 2014; García-Callejas & Araújo, 2015). Although the link was established based on spatially continuous predictions of Maxent outputs, the predictive accuracy was not assessed consistently and it was unclear whether optimal models selected with information criteria (AIC or BIC) tend to exhibit a high accuracy in predicting presence and absences sites. Our results were consistent across a series of validation metrics based on the confusion matrix and direct comparisons with true presence-absence models. Therefore, we suggest that model complexity measured through AIC is not related with accuracy in prediction-oriented ecological niche models.

Some authors have recommended against using a threshold criterion to generate binary predictions in ENMs (Merow et al., 2013). The criticism relies on the necessity to make strong assumptions about which threshold criteria use (Liu et al., 2005). However, this is not convenient because many statistical methods for fitting ENMs uses different functions to generate geographical predictions and therefore the continuous outputs are not necessarily comparable between them (Merow et al., 2014). We suggest that always will be better to thresholding continuous maps in binary predictions using a threshold value if we are interested in predict a potential or realized geographical distribution. We did it here and we found that results were consistent across several threshold criteria. Indeed, we found that almost all models predicted relatively well the independent presences dataset, independently of the regularization or feature used (Figures S10-S11). Finally, many models over-predicted the “true” geographic distribution (i.e., absence sites were identified incorrectly as presence sites). These results were consistent with previous studies showing that Maxent algorithm tends to extrapolate further “known” niche conditions (Saupe et al., 2014).

Our results support the idea that for users interested in prediction-oriented models is better to evaluate which regularization and feature used are better to predict the independent dataset of presences and absences (or pseudo-absences). If users are interested in explanation-oriented models (i.e., estimate niche conditions across geography, or establish what variables determine the presence-absence of a species), strategies to evaluate model complexity as AIC or BIC are warranted. However, we suggest that in that cases it might be necessary to evaluate the performance of different parameterization schemes explicitly in the ecological space. For instance, it is well-known that ecological niche (Grinnellian niche, Soberón, 2007; Peterson et al., 2011) is defined in terms of the environmental conditions that allow a population have an intrinsic positive per-capita growth (i.e., source populations; McGuire, 1973; Holt, 2009; Pulliam, 2000). Also, it is well-known that sink populations, which are only maintained from dispersals from other sources, are outside the niche of a species (Holt, 2009; Pulliam, 2000). Accordingly, it is crucial to evaluate whether optimal ecological niche models are able to discriminate between sink and source populations based on its preferred environmental conditions.

## Acknowledgments

We are grateful with Enrique Martínez Meyer, Angela Cuervo, Luis Osorio and members of the Laboratorio de Análisis Espaciales at IB-UNAM for discussions about this topic. We also grateful with Town Peterson, Luis Escobar, and Luis Osorio-Olvera for extensive and thoughtful comments to a previous version of this manuscript. Luis Osorio-Olvera also help us with R scripts to generate virtual species and validation. JAV is funded by a DGAPA postdoctoral scholarship at UNAM.

## Biosketch

**Julian A. Velasco** is a biologist interested in biogeography and macroecology of Neotropical taxa. He also is interested in spatially-explicit methods to generate geographical predictions of multiple biodiversity dimensions at different spatial scales.

**Constantino Gonzales-Salazar** is an ecologist interested in ecological niche modeling and prediction of diseases in geographical space using mining data approaches

